# Neural evidence for representationally specific prediction in language processing

**DOI:** 10.1101/243667

**Authors:** Lin Wang, Gina Kuperberg, Ole Jensen

## Abstract

Previous studies suggest that people generate predictions during language comprehension at multiple linguistic levels. It has been hypothesized that, under some circumstances, this can result in the pre-activation of specific lexico-semantic representations. We asked whether such representationally specific semantic pre-activation can be detected in the brain ahead of encountering bottom-up input. We measured MEG activity as participants read highly constraining sentences in which the final word could be predicted. We found that both spatial and temporal patterns of the brain activity prior to the onset of this word were more similar when the same words were predicted than when different words were predicted. This pre-activation was transient and engaged a left inferior and medial temporal region. These results suggest that unique spatial patterns of neural activity associated with the pre-activation of distributed semantic representations can be detected prior to the appearance of new sensory input, and that the left inferior and medial temporal regions may play a role in temporally binding such representations, giving rise to specific lexico-semantic predictions.

## 1. Introduction

After reading or hearing the sentence “In the crib there is a sleeping …”, the word “baby” is already activated in our minds before we encounter it. Here, we used magnetoencephalography (MEG) to determine whether it is possible to detect a spatio-temporal signature that corresponds to the representationally specific pre-activation of the predicted word (“baby”).

Language prediction allows us to rapidly understand what we read or hear by giving processing a head start. Within a hierarchical generative framework of language processing^1^, information at higher levels flows down the hierarchy to pre-activate information at lower levels in an attempt to minimize prediction errors within an internal generative model that explains the statistical structure of the bottom-up sensory input. When these predictions match this bottom-up input, its recognition is facilitated, and when they mismatch this input, the generative model is updated. During language processing, people can generate probabilistic predictions at multiple linguistic levels, including events^2^, syntax^3^, semantic features^4^, phonetic^5^ and orthographic features^6^ and even lower-level visual^7^ and auditory^8^ features. The level and strength of such pre-activation depends on many factors, including contextual constraint^9^ and the comprehender’s communicative goals and strategy (see ^1^, section 3.4).

Prediction is also hypothesized to be a core computational principle of brain function^10^. Numerous electrophysiological studies have investigated language prediction by measuring brain responses to words with different levels of predictability. This includes a wide range of studies based on the N400 paradigm^11^. Only a few studies, however, have examined brain activity associated with the prediction period itself, prior to the presentation of the predicted word. These studies report that highly constraining sentence contexts induce an increase in theta power^12,13^ as well as a suppression in alpha/beta power^14–17^. They also report activity within both neocortical (e.g., left frontal and temporal regions;^12,15^) and subcortical (e.g., hippocampus and cerebellum;^13,18,19^) regions. Thus far, however, no study has determined whether it is possible to detect neural activity associated with representationally specific lexico-semantic prediction, and, if so, when exactly such information is active in relation to the appearance of the bottom-up input.

Neural patterns reflecting representationally specific information can be detected using multivariate pattern analysis (MPVA) such as representational similarity analysis (RSA)^20–22^. RSA assumes that similarities in both spatial and/or temporal patterns of brain activity can be used to identify brain activity associated with representationally similar items. This approach has, for example, been used to decode representationally specific visual information during both perception^23,24^ and working memory maintenance^25,26^.

In the present MEG study, participants read pairs of sentences with highly constraining contexts (Fig. 1a). Each member of a pair predicted the same word. We visually presented the sentences at a slow rate of 1s per word. This guaranteed sufficient time to detect any representationally specific neural activity before the onset of the predicted word, and to identify its precise time course in relation to the appearance of the bottom-up input. We hypothesized that, if representationally specific lexico-semantic information is pre-activated, the brain activity associated with pre-activation of the same predicted words (*within-pairs*) should be more similar than brain activity associated with differently predicted words (*between-pairs*). We examined both spatial and temporal indices of neural similarity. Spatial similarity values were computed by correlating the pattern of MEG data across all sensors between pairs of sentences. This enabled us to identify the temporal dynamics associated with the pre-activation of specific lexico-semantic items (Fig. 1b). Temporal similarity values were computed by correlating the temporal pattern of MEG data between pairs of sentences. By examining such temporal correlations at each location (Fig. 1c), we were able to identify the key brain structures involved in generating representationally specific semantic predictions.

## 2. Results

Twenty-six participants read 120 sentence pairs presented at a rate of one word per second while MEG data were acquired. Pairs of sentences were constructed that strongly predicted the same sentence-final words (SFWs). As an example (Fig. 1a), sentences S1 (e.g. “In the hospital, there is a newborn…”) and S2 (“In the crib there is a sleeping…”) both predicted the word “baby”. We compared the similarity pattern of sentence pairs that predicted the same SFWs (*within-pairs*) to those that predicted different SFWs (*between-pairs*) before the SFW actually appeared. To avoid repetition of the predicted word across sentence pairs, one member of each pair ended with the predicted word (e.g. in S1, “baby”) while the other member ended with an unpredicted but plausible word (e.g. in S2, “child”). The 120 sentence pairs were presented in random order. Participants were asked to read each sentence carefully and to answer yes/no comprehension questions following 1/6th of the sentences. Comprehension accuracy was high (98%+/−2.0%).

### 2.1. The spatial pattern of neural activity was more similar in sentence pairs that predicted the same versus different words

In each participant, we quantified the degree of spatial similarity of MEG activity (30Hz low-pass filter) produced by pairs of sentences that predicted either the same SFW (i.e. *within-pair*, e.g. S1-A vs. S2-A’) or a different SFW (i.e. *between-pair*, e.g. S1-A vs. S3-B) by correlating the pattern of signal produced across sensors at each sampling point from -2000ms to 1000ms relative to SFW onset. We then averaged the resulting time series of spatial correlations (R-values) within each participant and then across participants (Fig. 2b). Both the *within*- and *between-pair* group-average time series showed a sharp increase at ~100ms after the onset of the second-from-final word (SFW-1; at -1000ms) that lasted ~400ms before sharply decreasing (Fig. 2a). The same pattern was observed around the previous word (SFW-2) and around the SFW. We attribute this general increase in spatial similarity to the visual onset and offset of each word, resulting in an increase of spatial similarity for around 300ms after its offset at 200ms relative to the word onset. We focused the rest of the analysis on the predictive intervals before the SFW and before SFW-1 where these general increases in spatial correlation were largest (R > 0.04).

Averaged across the −880 – −485ms interval before the onset of the SFW (corresponding to 120 – 515ms after the onset of SFW-1), we found that the spatial pattern of neural activity (across sensors) was more similar in sentence pairs that predicted the same SFW (*within-pairs*) than in pairs that predicted different SFWs (*between-pairs*), t_(25)_ = 4.434, p < .001, Fig. 2b. Fig. 2c shows a scatter plot of the averaged R-values per participant in this interval. Twenty-two out of 26 subjects had R-values below the diagonal, i.e. larger values for the *within-pair* than the *between-pair* spatial correlations. In contrast, there was no difference between the *within-pair* and *between-pair* spatial correlation values averaged across the −1900 – −1510ms interval before the SFW (corresponding to 100 – 490ms after the onset of SFW-2): t_(25)_ = −0.212, p = .834.

**Fig. 1.**
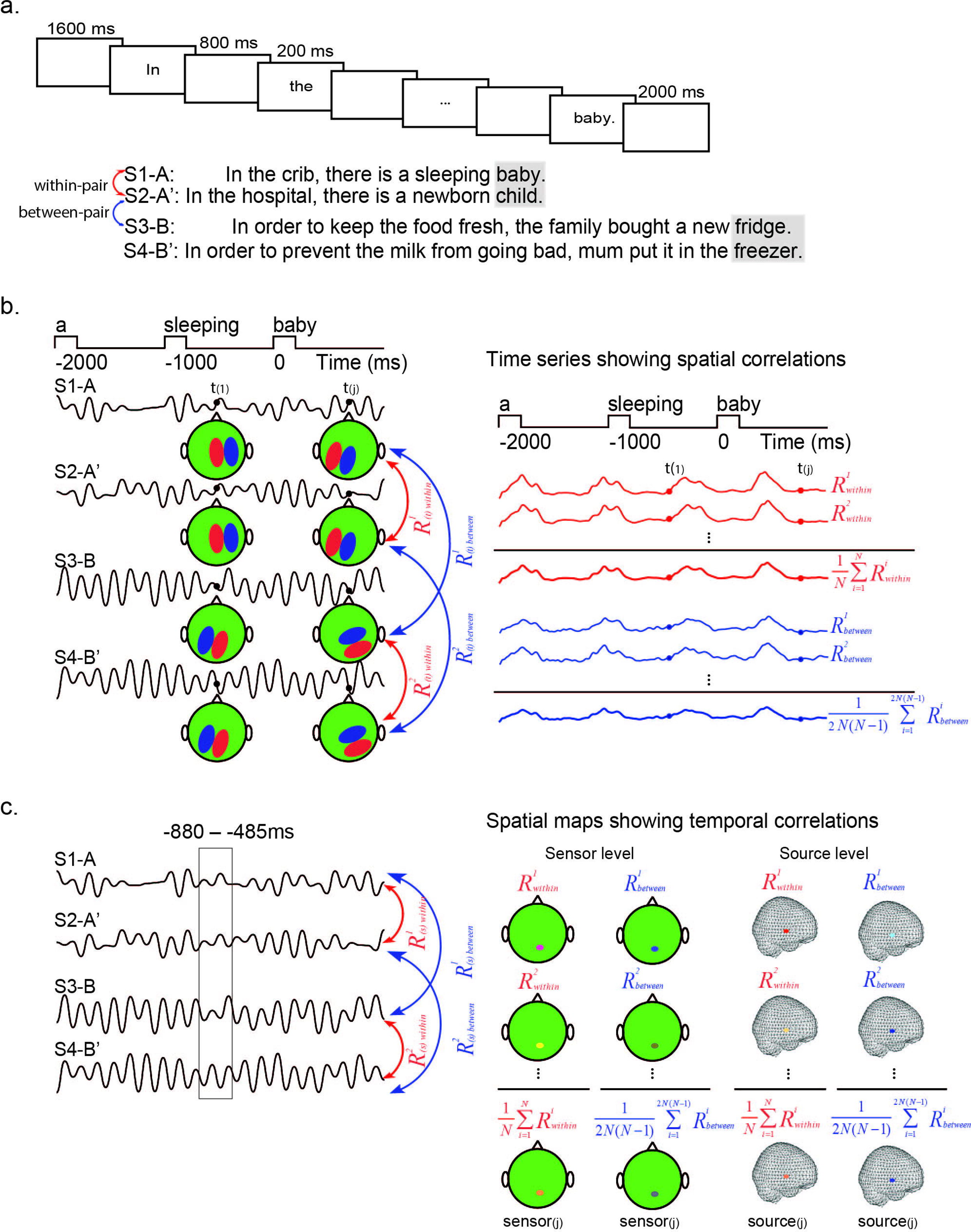
The experimental procedure and approach for Representational Similarity Analyses. (a) Trials began with a blank screen (1600ms). Sentences were presented in Chinese (translated here into English), word-by-word (200ms per word; 800ms blank interval between words). Sentences were followed either by ‘NEXT’ (2000ms) or by a probe question (1/6th of trials, randomly). We constructed sentence pairs in which the same word could be predicted from the context (e.g. S1-A & S2-A’; S3-B & S4-B’). One member of each pair ended with the predicted word (e.g. S1-A, S3-B) and the other member ended with a plausible but unpredicted word (e.g. S2-A’, S4-B’). Before the onset of the predicted word, we compared brain activity associated with the prediction of the same word (*within-pair*) and different words (*between-pair*). (b) Spatial Representational Similarity Analysis. Left: The pattern of MEG data over sensors was correlated between each sentence pair (*i* e.g. S1-A and S2-A’) at each time sample t_(j)._. Right: The average spatial correlation values for pairs 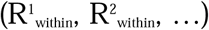 in which the same word was predicted formed the *within-pair* spatial correlation time series (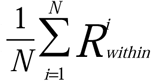, shown in red). The average of pairs 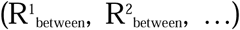 in which different words were predicted formed the *between-pair* spatial correlation time series (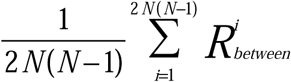, shown in blue). (c) Temporal Representational Similarity Analysis. Left: The pattern of MEG activity over time was correlated between sentence pairs, sensor-by-sensor or grid-by-grid point (source level). Right: The average temporal correlation values for pairs 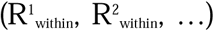 in which the same word was predicted formed the *within-pair* temporal correlation topographic/source maps. The average of pairs 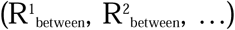 in which different words were predicted formed the *between-pair* temporal correlation topographic/source maps.

This difference in spatial similarity within the predictive interval cannot be explained by differences in the number of *within*- and *between-sentence* pairs used to compute these spatial correlation values because they were calculated per pair before averaging (see ^27,28^). However, to convince skeptics, we repeated the analysis using a randomly selected subset of *between-pair* correlations that matched the number of *within-pair* correlations. This analysis confirmed that the *within-pair* spatial correlation values were greater than the *between-pair* correlations (t_(25)_ = 2.393, p = .025; see Supplementary Fig. 1).

We avoided the repetition of the SFW (e.g. “baby”) between pairs by replacing the predicted SFW of one member with an unpredicted but plausible word (e.g. “child”). However, it might be argued that, after encountering the predicted word (“baby”), participants retained this item within memory and that the increased spatial similarity of brain activity when reading the other member of the pair was due to anticipatory retrieval facilitated by the previous presentation of the item. To address this concern, we divided the sentence pairs into two subsets according to whether the sentences with *expected* or *unexpected* SFWs were presented first. We then applied the spatial similarity analysis to both subsets (Supplementary Fig. 2) and compared their spatial similarity values. A repeated measures ANOVA with the factors Order (*Expected* SFW first, *Unexpected* SFW first) and Pairs (*Within-pair, Between-pair*) showed no main effect of Order (F_(1,25)_ = 0.747, p = .396, η2 = .029), nor an interaction between Order and Pairs (F_(1,25)_ = 1.804, p = .191, η2 = .067). We conclude that previously encountering a sentence ending with the expected SFW did not inflate the spatial similarity between sentence pairs that predicted the same SFW.

**Fig. 2.**
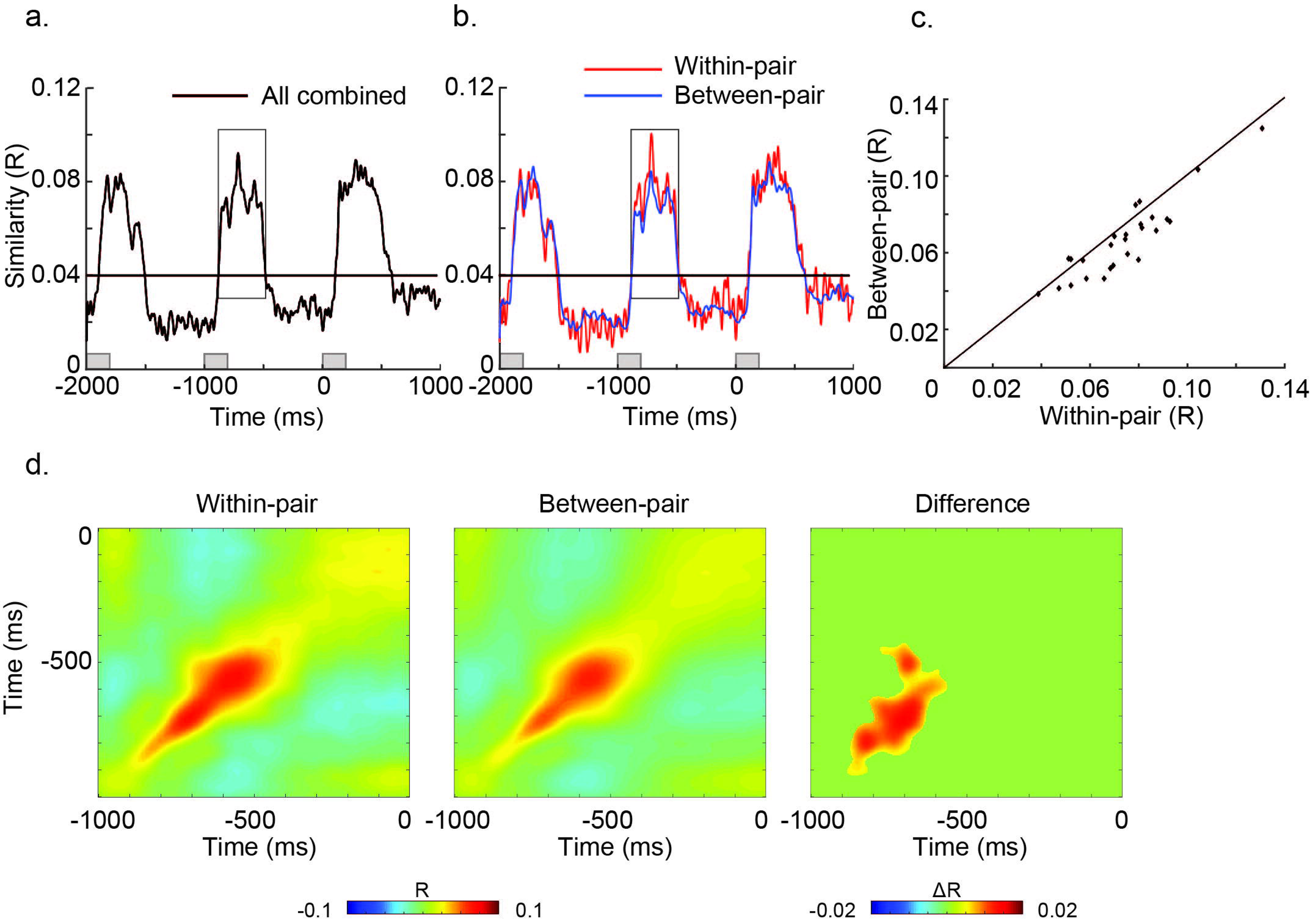
Results: Spatial Representational Similarity Analysis. (a) The time series of spatial similarity R values combined across *within-pair* and *between-pair* correlations, with the horizontal line indicating a threshold of R = 0.04 where the general increase in spatial correlation was largest. (b) The time series of spatial similarity R values for pairs in which the same word was predicted (*within-pair*, shown in red) and in which different words were predicted (*between-pair*, shown in blue). Both the *within*- and the *between-pair* spatial similarity time series showed a sharp increase at ~100ms and a decrease at ~400ms after the onset of each word. Between −880 – −485ms before the onset of the final word, the spatial similarity was greater when the same word was predicted than when different words were predicted (t(_25_) = 4.434, p < .001). (c) Scatter plots of spatial similarity values averaged between −880 – −485 ms before the onset of the final word in 26 participants. In most participants (22/26) the *within-pair* spatial correlations were greater than the *between-pair* spatial correlations. (d) Cross-temporal spatial similarity matrices for the *within*- and *between-pair* correlations (Red: positive correlations; blue: negative correlations). Left & middle: Both sets of pairs showed increased spatial similarity along the diagonal with greater similarities for the *within*- than the *between-pairs* in the −900 – −500ms interval prior to the onset of the final word. Right: The matrix shows the cluster with a statistically significant difference between the *within-pair* and *between-pair* spatial correlations (p = .002, cluster-randomization approach controlling for multiple comparisons over time by time samples). The absence of ‘off-diagonal’ correlations suggests that the spatial pattern associated with prediction was reliable but changed over time.

To characterize the temporal dynamics of brain activity reflecting representationally specific predictions, we correlated the spatial pattern of activity (across sensors) between one sentence (e.g. S1-A) at a particular time sample (e.g. t_1_) with that of its paired sentence (e.g. S2-A’) at all time samples (e.g. from t_1_ to t_n_) in each participant (see also ^29^ and ^21^), yielding a cross-temporal *within-pair* similarity matrix. We also calculated *between-pair* cross-temporal similarity matrices, and averaged them within each participant and then across participants (Fig. 2d). As expected, both the *within*- and *between-pair* group-averaged cross-temporal spatial similarity matrices showed that the spatial similarity was strongest around the diagonal in the first half-second after the onset of SFW-1. This was also the case for the difference between *within-pair* and the *between-pair* matrices (cluster-based permutation test: p = .002). This effect along the diagonal is consistent with the spatial similarity difference reported in Figs. 2b and 2c. Importantly, the absence of an effect off the diagonal suggests that the spatial patterns associated with prediction were not stable over time.

### 2.2. The temporal pattern of neural activity was more similar in sentence pairs that predicted the same versus different words, and this effect localized to left inferior temporal regions

As described above, across the −880 – −485ms interval prior to the onset of the SFW, we observed a general increase in spatial similarity between all pairs of sentences, regardless of whether they predicted for the same (*within-pair*) or a different SFW (*between-pair*). We next asked whether, within this time window, there were any brain regions in which the temporal pattern of neural activity was more similar between sentences that predicted the same versus different SFWs. To address this question, in each participant, we quantified the degree of temporal similarity of MEG activity produced by pairs of sentences that predicted the same versus different SFWs by correlating the temporal pattern of signal produced within this time window at each sensor. We then averaged the resulting spatial topographic maps of temporal correlations within each participant and then across participants (Fig. 1c). The group-averaged temporal similarity maps revealed a general increase in temporal similarity over bilateral temporal and posterior sensors, regardless of whether sentences predicted the same or a different SFW. When comparing the *within*- and *between-pair* temporal similarity topographic maps (Fig. 3a), the temporal pattern of neural activity was more similar in pairs that predicted the same versus different words over central and posterior sensors (cluster-based randomization test: p = .008; Fig. 3a: right panel).

In order to estimate the underlying neuroanatomical source of the increased temporal similarity associated with the *within-pair* sentences, we repeated this analysis in source space. We first discretized the full brain volume using a grid. At each grid point, we constructed spatial filters at each grid point using a ‘beamforming approach’ (a linearly constrained minimum variance technique^30^) and applied it to the MEG data. Then we performed the temporal similarity analysis on the time series from the spatial filters. The differences in the temporal similarity R-values were mapped on the grid in each participant. These difference values were then morphed to the MNI brain and averaged. This analysis showed that the temporal pattern of neural activity was more similar in sentence pairs that predicted the same versus different words within the left inferior temporal region (see Supplementary Fig. 4a for the 85% maximum difference). The source extended medially to include the left fusiform, parahippocampus and hippocampus, as well as the left cerebellum (Fig. 3b; cluster-randomization controlling for multiple comparisons: p = .004). The comparison of a randomly selected subset of *between-pair* correlations that matched the number of *within-pair* correlations confirmed this finding (cluster-randomization controlling for multiple comparisons: p = .034, see Supplementary Fig. 3b for the statistically significant cluster and Supplementary Fig. 4b for the 85% maximum difference).

**Fig. 3.**
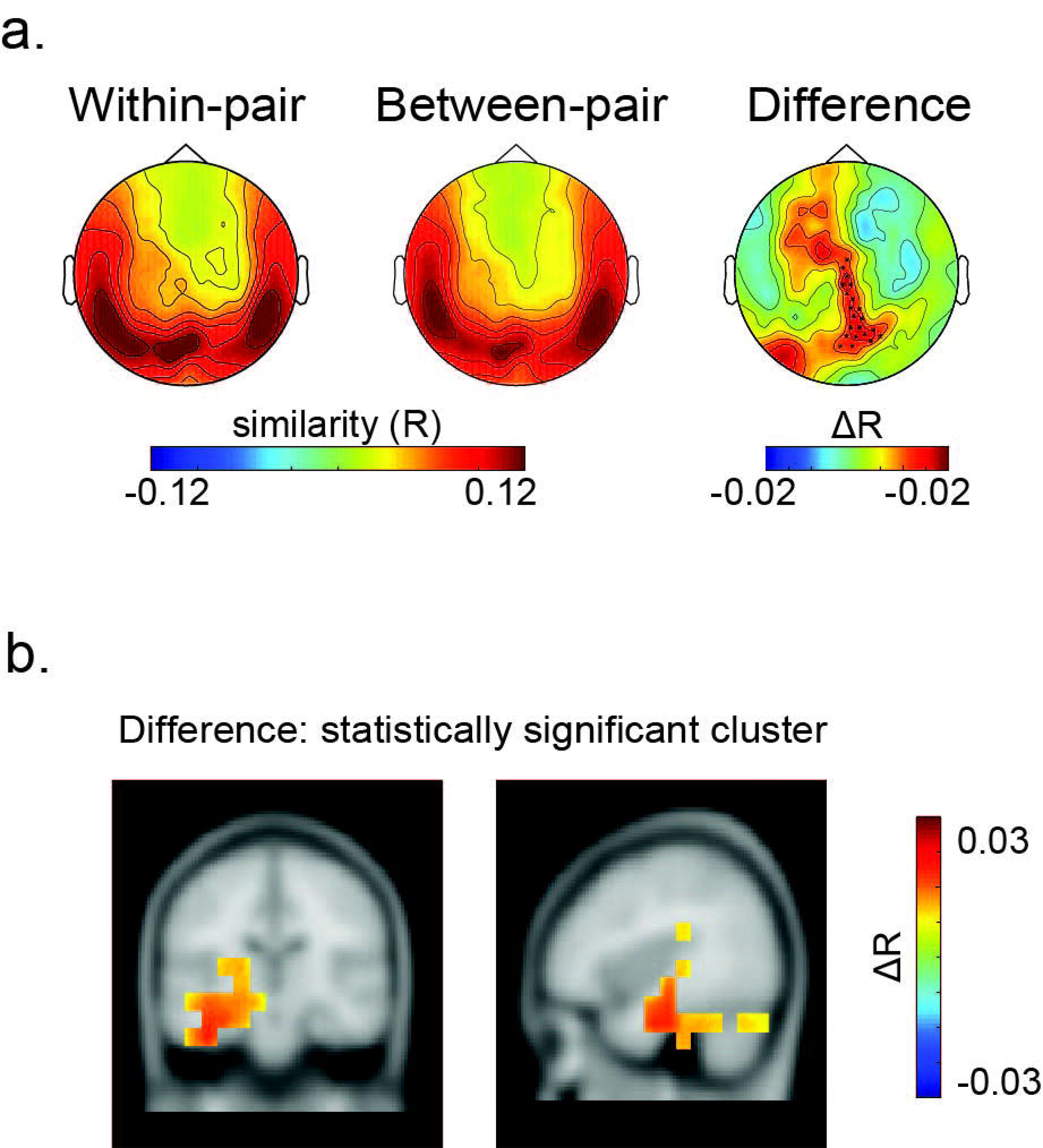
Results: Temporal Representational Similarity Analysis carried out between −880 – −485ms before the onset of the final word. (a) Temporal similarity topographic maps at the sensor level. Left & middle: Both the *within*- and *between-pair* correlations revealed increased temporal similarity over bilateral temporal and posterior sensors. Right: the difference map revealed greater temporal similarity when the same word was predicted than when different words were predicted over central and posterior sensors. The cluster of sensors where this difference was significant is marked with black asterisks (p = .008; a cluster-randomization approach controlling for multiple comparisons over sensors). (b) Temporal similarity difference map in source space. The values were interpolated on the MNI template brain and are shown both on the coronal plane (Talairach coordinate of peak: y = −19.5 mm) and the sagittal plane (Talairach coordinate of peak: x = −39.5 mm). This revealed significantly greater temporal similarity between sentence pairs that predicted the same word than pairs that predicted different words within the left inferior temporal gyrus, extending into the medial temporal lobe including the left fusiform, hippocampus and parahippocampus as well as left cerebellum (p = .004; a cluster-randomization approach controlling for multiple comparisons over grid points).

## 3. Discussion

We aimed to identify neural activity associated with representationally specific lexico-semantic prediction during sentence reading. To this end, we used MEG in conjunction with a representational similarity approach to index brain activity as participants read sentences in which the last word was highly predictable from the context. Based on a spatial correlation measure we were able to detect a unique pattern of spatial activity associated with representationally specific lexico-semantic prediction. This activity was evident between 120 and 515ms following the word before the predicted word (SFW-1). Moreover, within this predictive window, a temporal correlation measure implicated the left inferior temporal region and neighboring areas as playing an important role in implementing specific lexico-semantic prediction. To the best of our knowledge, this is the first study to detect representationally specific lexico-semantic pre-activation during language processing.

### 3.1. Representationally specific pre-activation appeared immediately after the prediction was available

Our finding that the spatial pattern of neural activity (across sensors) was more similar in sentence pairs that predicted the same SFW (*within-pairs*) than in pairs that predicted different SFWs (*between-pairs*) provides strong evidence that representationally specific information was pre-activated. We argue that the increased *within-pair* spatial correlations reflected the similarity between unique spatial patterns of neural activity across sensors associated with the same predicted word. We suggest that these unique spatial patterns corresponded to the pre-activation of a unique sets of distributed semantic features associated with the predicted words.

An alternative possibility is that, rather than reflecting pre-activation of the same word, the increased spatial similarity associated with the *within-pairs* reflected greater similarity between their *sentence contexts*. We think this is unlikely, since the two contexts within each pair were composed of distinct words; in particular, the word before the SFW (SFW-1) always differed within pairs. Further, the effect was first observed in the 120 – 515ms interval after the onset of word before the SFW (SFW-1). If the effect was driven by related contexts, the larger *within-pair* than *between-pair* correlations would have started to build up earlier. The spatial similarity effect also cannot be attributed to the simple recognition of a previously presented SFW (e.g. “baby”) in the within-pair analysis. This is because the spatial similarity effect was just as large when the unexpected SFW of a pair was presented before the expected SFW as in the opposite order (see Supplementary Fig. 2).

We deliberately presented the sentences at a slow rate of 1000ms per word, and measured spatial similarity values at each point in time in order to probe the precise time-course of any representationally specific pre-activation in relation to the appearance of the bottom-up input. The increased spatial similarity between sentence pairs that predicted the same (versus a different) SFW only began at around 120ms after the onset of the word before the SFW (SFW-1). This was the first point in time at which participants had sufficient information to unambiguously predict a specific SFW. This increased spatial similarity effect lasted until 515ms following the onset of the SFW-1, corresponding to 315ms following its offset and then dropped off in the second half of the interval before the onset of the SFW (see Fig. 2). The cross-temporal spatial similarity matrix (Fig. 2d) confirmed the dynamic nature of this effect. It suggests that, rather than being maintained by persistent neural activity across time, any representationally specific predictive activity was transient in nature. This finding is in line with the notion of ‘activity-silent’ working memory^31,32^. Of course, it will be important for future work to determine whether similar dynamics are associated with representationally specific lexico-semantic prediction when bottom-up inputs unfold at faster, more naturalistic rates.

### 3.2. Representationally specific pre-activation was associated with left inferior and medial temporal activity

By analyzing temporal similarity at the source level during the prediction period, we were able to identify regions that showed more similarity in their temporal pattern of neural activity in sentence pairs that predicted the same (*within-pair*) than different SFWs (*between-pair*). We found evidence of increased temporal similarity within the left inferior temporal gyrus, extending into the medial temporal lobe, including the left fusiform, and parahippocampal gyrus and the hippocampus. We suggest that these regions played a functional role in temporally binding the unique patterns of spatial activity (corresponding to unique combinations of distributed semantic features) that were pre-activated prior to the onset of the SFW, and therefore in instantiating representationally specific lexico-semantic prediction.

The left inferior temporal lobe, particularly its more anterior portions, has long been implicated in lexico-semantic processing^33–39^. In particular, it has been proposed that it acts as a hub (a “convergence zone”^40^) that brings together conceptual information that is distributed throughout cortex^41,42^. Parts of this region may play a particular role in lexical processing, functioning as the brain’s dictionary by mediating between conceptual and phonological knowledge of words^43,44^. Ventral anterior subregions of the temporal cortex have also been implicated more generally in subserving not only verbal but also non-verbal multimodal semantic representations^41,45–47^. Most relevant to the present study is a recent finding that temporal RSA applied to intracranial EEG signals was associated with neural activity within the left inferior temporal lobe that encoded item-specific representations during picture naming^24^. Together with the present findings, this supports the idea that the temporal pattern of activity within this region plays a functional role in instantiating the activation of specific lexico-semantic items. For example, by tracking the precise time-course of activity within distributed regions, it may act to bind them together to form a coherent whole^48^. Crucially, in present study, the increase in temporal similarity between sentences that predicted the same SFW began before the bottom-up input became available, indicating that it may also instantiate representationally specific lexico-semantic prediction.

Of potential relevance to the idea that this region plays a role in temporal binding is the observation that it extended medially to include the hippocampus. While MEG source-modeling results within medial and subcortical regions should be interpreted with caution, the possible involvement of the hippocampus is very interesting given other work that has implicated it as playing a crucial role in binding representations to generate predictions. A large literature from recordings in rats demonstrates that the hippocampus represents upcoming spatial representations as the rat is navigating^49^, and we have a good understanding of the physiological mechanisms supporting such predictions^50^. There is also growing evidence that these predictive mechanisms might generalize to the human hippocampus^51–55^. Moreover, recently, it was found that the temporal patterns in higher frequency bands recorded within the hippocampus were similar between a pre-picture interval and the picture itself^56^, suggesting a role in representing pre-activated non-verbal semantic information. Given these findings, it is conceivable that the hippocampus also plays an analogous role in language prediction. Indeed Piai et al. (2016) used intracranial recordings in humans to demonstrate predictive effects in the hippocampus in a language task in which the sentence-final word had to be produced^13^.

In conclusion, we successfully used MEG to identify unique spatial and temporal patterns associated with the prediction of specific lexico-semantic items during language processing. We showed that the spatially specific neural patterns became active at around 100ms after a word was unambiguously predicted, that this pre-activation was transient and that it was accompanied by unique temporal patterns of activity within the left inferior and medial temporal lobe. These findings pave the way towards the use of these methods to determine whether and when such specific lexico-semantic representations are pre-activated as language, in both visual and auditory domains, unfolds more rapidly in real time.

## Acknowledgements

This work was funded by the Natural Science Foundation of China (31540079 to LW), the National Institute of Child Health and Human Development (R01 HD08252 to GRK), and a James S. McDonnell Foundation Understanding Human Cognition Collaborative Award (220020448) and the Royal Society Wolfson Research Merit Award to OJ. It was supported in part by the Ministry of Science and Technology of China grants [2012CB825500, 2015CB351701]. We thank Yang Cao and Yinan Hu for their assistance with data collection.

---

***Supplementary Fig. 1.*** Results: Spatial Representational Similarity Analysis after matching the number of pairs between the *within-pair* and *between-pair* correlations. (a) The time series of spatial similarity R values for the pairs in which the same word was predicted (*within-pair*, shown in red) and in which a different word was predicted (*between-pair*, shown in blue). Within the −880 – −485ms interval relative to the onset of the final word, the spatial similarity was greater when the same word was predicted than when different words were predicted (−880 – −485ms before its onset; t_(25)_ = 2.393, p = .025). (b) Scatter plots of the spatial similarity values averaged between −880 – −485ms before the onset of final word in 26 participants. In most participants (17/26) the *within-pair* spatial correlations were greater than the *between-pair* spatial correlations.

***Supplementary Fig. 2.*** Results: Spatial Representational Similarity Analysis for two subsets of trials where sentences ending with *expected* words were seen first (a and b) or sentences ending with *unexpected* words were seen first (c and d). The time series of spatial similarity R values for the pairs in which the same word was predicted (*within-pair*) are shown in red, while the time series for the pairs in which a different word was predicted (*between-pair*) are shown in blue. The spatial similarity was greater when the same word was predicted than when different words were predicted in both subsets. No significant difference was found between the two subsets of trials, as indicated by the lack of a main effect of Order (*Expected* First, *Unexpected* First) (F_(1,25)_ = .747, p = .396, η2 = .029) or an interaction between Order (Expected First, Unexpected First) and Pairs (*Within-pair, Between-pair*) (F_(1,25)_ = 1.804, p = .191, η2 = .067).

***Supplementary Fig. 3.*** Results: Temporal Representational Similarity Analysis carried out between −880 – −485ms before the onset of the final word after matching the number of pairs between the *within-pair* and *between-pair* correlations. (a) Temporal similarity topographic maps at the sensor level. Left & middle: Both the *within*- and *between-pair* correlations revealed increased temporal similarity over bilateral temporal and posterior sensors. Right: the difference map revealed greater temporal similarity when the same word was predicted than when different words were predicted over central and posterior sensors (marginally significant cluster: p = .0679; a cluster-randomization approach controlling for multiple comparisons over sensors). (b) Temporal similarity difference map in source space. The values were interpolated on the MNI template brain and are shown both on the coronal plane (Talairach coordinate of peak: y = −9.5 mm) and the sagittal plane (Talairach coordinate of peak: x = −29.5 mm). This revealed significantly greater temporal similarity between sentence pairs that predicted the same word than pairs that predicted a different word within the left hippocampus and extended into the left inferior temporal region, left fusiform, parahippocampus, amygdala as well as left cerebellum (p = .034; a cluster-randomization approach controlling for multiple comparisons over grid points).

***Supplementary Fig. 4.*** Results: Temporal Representational Similarity Analysis carried out between −880 – −485ms before the onset of the final word, showing the 85% maximum difference of the statistically significant cluster in source space. (a) Temporal similarity difference map in source space between the averaged N *within-pair* correlations and 2N(N-1) *between-pair* correlations. The values were interpolated on the MNI template brain and are shown both on the coronal plane (Talairach coordinate of peak: y = −19.5 mm) and the sagittal plane (Talairach coordinate of peak: x = −39.5 mm). The maximum difference between the *within-pair* and the *between-pair* correlations was found within the left inferior temporal gyrus, and the cluster extended into the medial temporal lobe including the left fusiform, hippocampus and parahippocampus. (b) Temporal similarity difference map in source space between the averaged N *within-pair* correlations and N *between-pair* correlations. The values were interpolated on the MNI template brain and are shown both on the coronal plane (Talairach coordinate of peak: y = −9.5 mm) and the sagittal plane (Talairach coordinate of peak: x = −29.5 mm). The maximum difference between the *within-pair* and the *between-pair* correlations was found within the left inferior temporal gyrus, and the cluster extended into the medial temporal lobe including the left fusiform, hippocampus and parahippocampus.

## 4. Methods

### 4.1. Design and development of stimuli

We developed a stimulus set of 120 pairs of sentences in Mandarin with highly constraining contexts. The two contexts within each pair were distinct from one another, and had no content words in common (with the exception of five pairs), but they each strongly predicted the same sentence-final word (SFW). For example, e.g. in Fig. 1, both sentences S1 and S2 predicted the word, “baby”. In half of these sentences, the expected final word was a noun and in the other half, it was a verb.

To select and characterize this final set of sentences, we began with an initial set of 208 pairs and carried out a cloze norming study in 30 participants (mean age: 23 years; range: 18 – 28 years old; 15 males), who did not participate in the subsequent MEG study. In this study, sentence contexts were presented without the SFW (e.g. ‘In the crib there is a sleeping …’) and participants were asked to complete the unfinished sentence by writing down the most likely ending. The two members of each sentence pair were counterbalanced across two lists (with order randomized within lists), which were each seen by half the participants. Testing took approximately 40 minutes per subject.

To calculate the lexico-semantic constraint of each sentence context, we tallied the number of participants who produced the most common completion for a given context. We retained 66 pairs in which 73% of the participants predicted the same SFW, i.e. at least 11 out of 15 participants filled in the same word in each sentence pair. We then revised 103 sentences (54 sentences in list 1 and 49 in list 2) to make them more constraining, and we re-tested them in the same group of participants. After this second round of cloze testing, we selected the final set of 120 sentences for the MEG experiment. In the final set of stimuli, the lexico-semantic constraints of 109 pairs were above 70% and the constraints of the remaining 12 pairs were slightly lower (Mean: 58%; SD: 12). Across all pairs, the mean lexico-semantic constraint was 88% (SD: 12).

We then generated full sentences by adding a SFW to each member of a pair. In one member of the pair, this SFW was highly predictable; it was the most common word filled by the cloze participants (e.g. “baby” following context S1, “In the crib there is a sleeping…”). In the other member of the pair, we selected a word that was semantically related to the highly predicted word but was not produced by any of the participants in the cloze norming, with the whole sentence still being plausible (e.g. “child” following context, S2, “In the hospital, there is a newborn…”). Thus, for this sentence, the lexical cloze probability was zero, see Fig. 1a for examples. All sentence contexts (e.g. S1 and S2) were combined with both lexically predicted (e.g. A: ‘baby’) and unpredicted (e.g. A’: ‘child’) SFWs, e.g. S1-A, S1-A’, S2-A, S2-A’. All combinations of the sentence contexts and SFWs were then counterbalanced across two lists. This ensured that, in the MEG session, while each participant would see both members of each sentence pair, they never saw the same SFW twice. Within each list, sentences were pseudo-randomized so that participants did not encounter more than three expected or unexpected SFWs in succession. All Mandarin sentences, together with their English translations, are available in the Supplementary Material.

### 4.2. Participants in the MEG study

The study was approved by the Institutional Review Board (IRB) of the Institute of Psychology, Chinese Academy of Sciences. Thirty-four students from the Beijing area were initially recruited by advertisement. They were all right-handed native Chinese speakers without histories of language or neurological impairments. All gave informed consent and were paid for their time. The data of eight participants were subsequently excluded because of technical problems, leaving a final MEG dataset of 26 participants (mean age 23 years, range 20 – 29; 13 males).

### 4.3. Procedure

MEG data were collected while participants sat in a comfortable chair within a dimly-lit shielded room. Stimuli were presented on a projection screen in grey color on a black background (visual angle ranging from 1.22 to 2.44 degrees). As shown in Fig. 1a, each trial began with a blank screen (1600ms), followed by each word with an SOA of 1000ms (200ms presentation with an inter-stimulus interval, ISI, of 800ms). The final word ended with a period followed by a 2000ms ISI. After one-sixth of the trials, at random, participants read either a correct or an incorrect statement that referred back to the semantic content of the sentence that they just read (for example, S1-A and S2-A’ in Fig. 1a might be followed by the incorrect statement, “There is an old man.”). Participants were instructed to judge whether or not the statements were correct by pressing one of two buttons with their left hand. This helped ensure that participants read the sentences for comprehension. In all other trials, the Chinese word ‘ ▪ ▪ ’ (meaning ‘NEXT’) appeared, and participants were instructed to simply press another button with their left hand within 5000ms in order to proceed to the next trial.

The 240 sentences were divided into 8 blocks, with each block lasting about 8 minutes. Between blocks there was a small break during which participants were told that they could relax and blink, but to keep the position of their heads still. Participants could start the next block by informing the experimenter verbally. The whole experiment lasted about 1.5 hours, including preparation, instructions and a short practice session consisting of eight sentences.

### 4.4. MEG data acquisition

MEG data was collected using a CTF Omega System with 275 axial gradiometers at Institute of Biophysics, Chinese Academy of Sciences. Six sensors (MLF31, MRC41, MRF32, MRF56, MRT16, MRF24) were non-functional and were therefore excluded from the recordings. The ongoing MEG signals were low-pass filtered at 300Hz and digitized at 1200Hz. Head position, with respect to the sensor array, was monitored continuously with three coils placed at anatomical landmarks (fiducials) on the head (forehead, left and right cheekbones). The total movement across the whole experiment was, on average, 8mm across all participants. In addition, structural Magnetic Resonance Images (MRIs) of 24 participants were obtained using a 3.0T Siemens system. During MRI scanning, markers were attached in the same position as the head coils, allowing for later alignment between these MRIs and the MEG coordinate system.

### 4.5. MEG data processing

MEG data were analyzed using the Fieldtrip software package, an open-source Matlab toolbox^57^. In order to minimize environmental noise, we applied third order synthetic gradiometer correction during preprocessing. Then, the MEG data were segmented into 4000ms epochs, time-locked from the onset of two words before the SFW (SFW-2) until 2000ms after the onset of the SFW. Within this 4000ms epoch, trials contaminated with muscle or MEG jump artifacts were identified and removed using a semi-automatic routine. After that, we carried out an Independent Component Analysis (ICA;^58,59^) and removed components associated with the eye-movement and cardiac activity from the MEG signal (about 5 components per subject). Finally, we inspected the data visually and removed any remaining artifacts. On average 96% +/− 3.4%) of trials were retained.

#### 4.5.1. Spatial Representational Similarity Analysis

##### 4.5.1.1. Calculation of spatial similarity time series

A schematic illustration of the spatial representational similarity analysis (RSA) approach is shown in Fig. 1b. First, we detrended and applied a 30Hz low pass filter to the MEG data. Next, in each participant, for each trial, and at each time sample, we extracted a vector of MEG data that represented the spatial pattern of activity across all 269 MEG sensors (6 of 275 sensors were not operational). We then quantified the degree of spatial similarity of MEG activity produced by the two members of each sentence pair predicting the same SFW (e.g. between S1-A and S2-A’, in Fig. 1a) by correlating the spatial vectors between each member of a pair at consecutive time sample across the 4000ms epoch. This yielded a time-series of correlations (Pearson R-values) reflecting the degree of spatial similarity at each time sample between sentences that predicted the same SFW (e.g. time-series 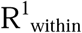 and 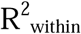, see Fig. 1b). We refer to these as *within-pair* spatial similarities. After artifact rejection, in each participant, there were, on average, N = 111+/−8 complete *within-pair* spatial similarity time series. We then averaged these time series together over all pairs of sentences that predicted the same SFW to yield an average *within-pair* spatial similarity time series within each participant (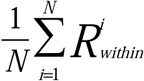; Fig. 1b).

We then repeated this entire procedure, but this time correlating spatial patterns of MEG activity between pairs of sentences that predicted a different SFW, for example, between S1-A and S3-B (Fig. 1a). This yielded 2N(N-1) *between-pair* spatial correlation time courses, e.g. 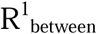 and 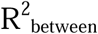 (Fig. 1b). We again averaged these together to yield a time series of R-values within each participant (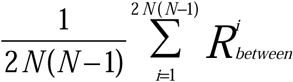; Fig. 1b), which reflected the degree of similarity between spatial patterns of activity elicited by sentences that predicted different SFWs at each time sample (i.e. *between-pair* spatial similarity time series). Fig. 2b shows the averages, across all participants, of the *within-pair* spatial similarity time series, and the *between-pair* spatial similarity time series.

##### 4.5.1.2. Calculation of cross-temporal spatial similarity matrices

To characterize how temporally sustained the spatial patterns were (see also^21,29^), we correlated the spatial pattern vector between one member of a sentence pair (e.g. S1-A) at a particular time sample (e.g. t_1_) with that of the other member of the pair (e.g. S2-A’) at all time samples (e.g. from t_1_ to t_n_) in each participant. The resulting values can be visualized as a similarity matrix (Fig. 2d), with each entry representing the spatial similarity between two sentences at two time samples (e.g. R_(i,j)_ represents the correlation between S1-A at time *i* and S2-A’ at time *j*). The R-values along the diagonal reflect the spatial similarity at corresponding time samples (R_(i,j)_ when i = j; i.e. the time of similarity R-values as described in Fig. 1b), while the R-values off the diagonal reflects cross-temporal spatial similarity. We repeated this procedure for pairs of sentences that predicted different SFWs (*between-pairs*). The cross-temporal similarity matrices for both *within-pair* and *between-pair* correlations, were averaged across sentence pairs for each participant (see Fig. 2d). Fig. 3 also shows the group-averaged *within-pair* and *between-pair* cross-temporal spatial similarity matrices. When constructing these cross-temporal similarity matrices, to increase computational efficiency, we down-sampled the data to 300 Hz and randomly selected N *between-pairs* to match with the N *within-pairs* for averaging. The correlation values were smoothed in time in both directions with a Gaussian kernel (40ms time window, SD: 8ms).

##### 4.5.1.3. Statistical testing

As can be seen in Fig. 2a, the averaged *within-pair* and the *between-pair* spatial similarity time series showed a sharp increase in R-values at around 100ms after the onset of the word before the SFW (SFW-1) lasting for about 400ms (i.e. 300ms into the ISI after the SFW-1 offset) before sharply decreasing again. This pattern of a sharp increase and decrease in spatial correlations was also seen in association with the previous word (SFW-2) as well as the following word (SFW). In order to objectively quantify the time-window over which this general increase in spatial similarity R values was sustained during the prediction period, we compared the averaged *within-pair* and *between-pair* spatial similarity time series against a threshold of R = 0.04 based on visual inspection of the R-values in the prediction time window. We found an increase in R-values from -880ms to -485ms (i.e. 120ms to 515ms relative to the onset of SFW-1), as well as from −1900 to −1510ms before the onset of the SFW (i.e. 100ms to 490ms relative to the onset of SFW-2) (Fig. 2b).

We then averaged across the −880 – −485ms interval before the onset of the SFW and carried out paired t-tests to determine whether, collapsed across these time windows, the spatial pattern of MEG activity produced by sentence pairs that predicted the same SFW was significantly more similar than the spatial pattern of MEG activity produced by sentences that predicted different SFW (i.e. *within-pair* vs. *between-pair* spatial correlation R values). We repeated the same analysis for the −1900 – −1510ms interval before the onset of the SFW

To test for cross-temporal statistical differences in spatial similarity patterns produced by sentence pairs that predicted the same versus different SFWs while controlling for multiple comparisons over time, we applied a cluster-randomization approach^60^. To this end, we first carried out paired t-tests at each data time sample in the cross-temporal spatial similarity matrices within the 1000ms interval between the onset of SFW-1 and the onset of SFW. We used temporal cluster-based permutations to account for multiple comparisons^60^. Data points that exceeded a pre-set uncorrected p-value of 0.05 or less were considered temporal clusters. The individual t-statistics within each cluster were summed to yield a cluster-level test statistic — the cluster mass statistic. We then randomly re-assigned the spatial similarity R values across the two conditions (i.e. *within-pair* and *between-pair*) at each data point within the matrix, within each participant, and calculated cluster-level statistics as described above. This was repeated 1000 times. For each randomization, we took the largest cluster mass statistic (i.e. the summed T values), and, in this way, created a null distribution for the cluster mass statistic. We then compared our observed cluster-level test statistic against this null distribution. Any temporal clusters falling within the highest or lowest 2.5% of the distribution were considered significant.

#### 4.5.2. Temporal representational similarity analysis

##### 4.5.2.1. Construction of temporal similarity maps at sensor level

In each participant, at each sensor for each trial, we considered the MEG time series in the −880 – −485ms interval before the onset of the SFW — that is, the time-window over which we observed the general increase in spatial similarity R values during the prediction period (see Fig. 2a). At each sensor, we then correlated this time series within this window between the two members of each sentence pair predicting the same SFW (e.g. between S1-A and S2-A’, in Fig. 1a) to yield an R value representing the degree of temporal similarity: an R value of 0 implies that the two time series are not correlated, while an R value of −1 implies that the two time series are anti-correlated. Together, these R values at each sensor yielded *within-pair* temporal similarity topographic maps for each pair, e.g. topographic maps 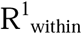 and 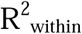, see Fig. 1c. We then averaged across all the *within-pair* temporal correlations at each sensor to yield an average *within-pair* temporal similarity topographic map within each participant and then averaged across participants (see Fig. 1c).

We then repeated this procedure, but this time correlating time series from MEG sensors produced by sentence pair that predicted different SFWs to investigate the spatial distribution of *between-pair* temporal correlations (e.g. topographic maps 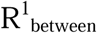 and 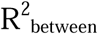 in Fig. 1c). These were again averaged together to yield an average topographic map of R values within participant, and then averaged across participants (Fig. 1c).

##### 4.5.2.2. Construction of temporal similarity maps at source level

We also constructed temporal similarity maps at the source level. We estimated the MEG signals at the source level by applying a spatial filter at each grid point using a beamforming approach^30^. We computed a linearly constrained minimum variance (LCMV; ^30^) spatial filter on the 30 Hz low-pass filtered (and linearly detrended) data from onset of SFW-1 to 1000ms after onset of SFW (i.e. −1000 – 1000ms relative to SFW onset). The LCMV approach estimates a spatial filter from a lead field matrix and the covariance matrix of the data from the axial gradiometers. To obtain the lead field for each participant, we first spatially co-registered the individual anatomical MRIs to the sensor MEG data by identifying the fiducials at the forehead and the two cheekbones. Then a realistically shaped single-shell head model was constructed based on the segmented anatomical MRI for each participant^61^. Each brain volume was divided into a grid with 10mm spacing and the lead field was calculated for each grid point. Then the grid was warped to the template Montreal Neurological Institute (MNI) brain (Montreal, Quebec, Canada). The MNI template brain was used for one participant whose MRI image was not available. The application of the LCMV spatial filter to the sensor-level data resulted in single-trial estimates of time series at each grid point in three orthogonal orientations. To obtain one signal per grid point we projected the time series along the direction that explains most variance using singular value decomposition. In order to construct temporal similarity maps in source space, we followed the same procedures as above, by correlating the time series at each grid point. The grand-average similarity values were interpolated onto the MNI template brain (Fig. 3b).

##### 4.5.2.3. Testing for significant difference between the within- and between-pair temporal similarity maps

To compare the *within-pair* vs. *between-pair* temporal correlation R value statistically, both at the sensor level and at the source level, we carried out a cluster-based permutation approach to control for multiple comparisons over sensors or grid-points^60^. At each sensor/grid point, in each participant, we compared the mean differences in the temporal similarity R values between sentence pairs predicting the same word (i.e. *within-pair*) versus a different word (i.e. *between-pair*). Sensors within 40mm that exceeded the 95^th^ percentile of the mean difference were considered clusters. In source space, clusters were formed by contiguous grids points. Within each cluster, we then summed the mean differences of R values at each sensor/grid-point to yield a cluster-level test statistic — the cluster mass statistic. Next, we randomly re-assigned the R-values across the two conditions (i.e. *within-pair* and *between-pair*) at each sensor/grid within each participant, and calculated cluster-level statistics as described above. This was repeated 1000 times. For each randomization, we considered the largest cluster mass statistic (i.e. the summed mean difference within a cluster to create a null distribution for the cluster mass statistic). Then we compared our observed cluster-level test statistic against this null distribution. Any clusters falling within the highest or lowest Design and development 2.5% of the distribution were considered significant.

